# Novel Electrodes for Reliable EEG Recordings on Coarse and Curly Hair

**DOI:** 10.1101/2020.02.26.965202

**Authors:** Arnelle Etienne, Tarana Laroia, Harper Weigle, Amber Afelin, Shawn K Kelly, Ashwati Krishnan, Pulkit Grover

## Abstract

EEG is a powerful and affordable brain sensing and imaging tool used extensively for the diagnosis of neurological disorders (e.g. epilepsy), brain computer interfacing, and basic neuroscience. Unfortunately, most EEG electrodes and systems are not designed to accommodate coarse and curly hair common in individuals of African descent. This can lead to poor quality data that might be discarded in scientific studies after recording from a broader population set, and for clinical diagnoses, lead to an uncomfortable and/or emotionally taxing experience, and, in the worst cases, misdiagnosis. In this work, we design a system to explicitly accommodate coarse and curly hair, and demonstrate that, across time, our electrodes, in conjunction with appropriate braiding, attain substantially (~10x) lower impedance than state-of-the-art systems. This builds on our prior work that demonstrated that braiding hair in patterns consistent with the clinical standard 10-20 arrangement leads to improved impedance with existing systems.

## I. Introduction

EEG is an inexpensive and noninvasive way to capture valuable information about an individual’s brain activity. EEG is the gold standard for epilepsy diagnosis [1], one of the first diagnostic steps for diagnosis of many neural disorders [2], [3], and one of the most common ways of brain-machine interfacing [4]. *Despite its widespread use, it is surprising, and extremely unfortunate, that the currently accepted process of recording EEG signals is not inclusive of every individual’s needs (equally surprising and unfortunate is the fact that this seems to have not been noted, much less addressed, in the academic community).* Specifically, existing EEG systems do not work well for patients with coarse, curly hair. Our work [5] was the first scientific study that observed the difficulties of applying EEGs to African hair. We discovered the problem through extensive discussions with (i) EEG technicians and neurologists, including Dr. Arun Antony and Dr. Christina Patterson, epilepsy neurologists at the University of Pittsburgh, (ii) researchers who extensively record EEG across the US^†^, and (iii) our own extensive experience on recording on individuals with coarse and curly hair, including the data reported in this paper. In this work, we propose a novel solution that combines braiding and novel electrode designs to record EEG from locations consistent with the clinical standard “10-20” recording system [7]. The goal of this work is as much to bring this problem to the attention of the engineering community, as it is to report data on our proposed novel solutions.

The core reason why African hair is unique comes down to follicle shape. Caucasian and Asian hair share many similarities: the follicle is more circular and large, which causes the hair to grow straight. In contrast, African hair comes from a smaller, more elliptical and flat follicles [9], [10]. Due to the shape of the follicle, as the hair grows, it curls (Fig. 1), causing difficulties in installation of EEG systems. We focus on “4A, 4B, and 4C” hair types (details on these hair-types and how they influence EEG are discussed in Appendix A).

**Fig. 1:**
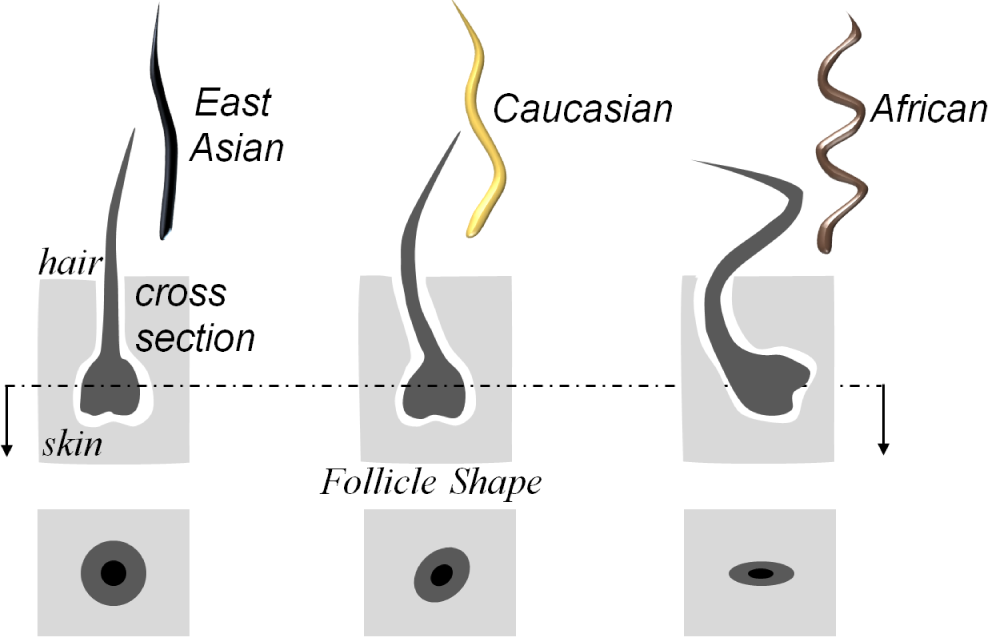
People of different ethnic origin have varying hair types that originate from different follicle sizes and shapes. People of African descent have particularly coarse and curly hair [8].

## II. Proposed Idea

In [5], we brought together fashion and technology to offer a simple solution to this problem: modifications on a styling method called *cornrowing*. Cornrowing is similar to braiding, but fashions the hair in a way that causes it to cling to the scalp and stay in place for long periods of time. It also creates exposed areas of skin (see Fig. 2) that are fixed and do not change with moisture and/or skin-abrasion (which is often used for applying electrodes). Cornrows can be worn for a long time (a few weeks). We modified traditional cornrowing to make it consistent with the 10-20 electrode positioning. In doing so, we adopted a simple style of cornrowing commonly referred to as “straight backs” (see Fig. 2). By comparing clinical standard gold cup electrodes on braided vs unbraided hair on a single participant, we observed in [5] that this simple solution can offer significant reductions in electrode-skin impedance. In this work, we report more data for this solution (totaling to 8 participants), as well as observe that, simply braiding the hair reduces the impedance, but the resulting impedance is still quite high (~ 2.4 MΩ at low frequencies). In this work, we develop novel electrodes for EEG, that we call *Sevo electrodes*^*2*^, that reduce impedance over gold cup electrodes *even for braided hair* by a factor of 10x.

**Fig. 2.**
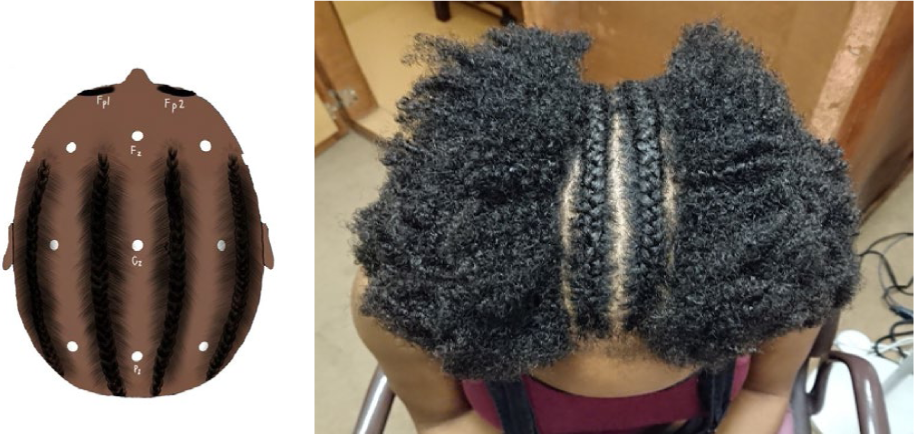
*(left)* A schematic of 10-20 consistent “straight-back” cornrowing. *(right)* Demonstration of cornrowing on a participant. The hair is braided down the head, exposing the scalp for electrode placement in locations consistent with 10-20 arrangement.

In this work, taking a step further, we address the issue of electrode design with coarse and curly hair. Currently, EEG systems are applied to the scalp of a patient in two ways: using a cap with electrodes embedded into it for higher density EEG, and small individual electrodes that are applied to strategic areas of the scalp that correspond to the desired brain region, which conventionally are gold cup electrodes. However, stretching a self-contained electrode cap over a large amount of hair or a relatively small amount of thick hair renders the electrode cap useless, as there is little to no contact between the system and the patient’s scalp. Additionally, even if the cap can reach the scalp, the electrodes often have minimal contact with the scalp, with the hair adding noise to and impeding the signal. This noise degrades the quality of data that can be obtained, hurting scientific studies and clinical inferences. The gold cup electrodes also pose a challenge as they are applied with a conductive paste that adheres the gold cup to the skin. Hair on the scalp can easily interfere with the electrode, and thus, the signal. This is especially true in coarse, curly hair types common in individuals of African descent.

To address these issues, we have designed an electrode-bearing clip (see Fig. 3) compatible for patients with coarse and curly that obviates the need for the electrode-bearing cap. This allows the EEG system to be entirely comprised of small independent electrodes that come together to form the conventional EEG system, such that the popular EEG configuration of 10-20 electrode placement can be preserved.

**Fig. 3:**
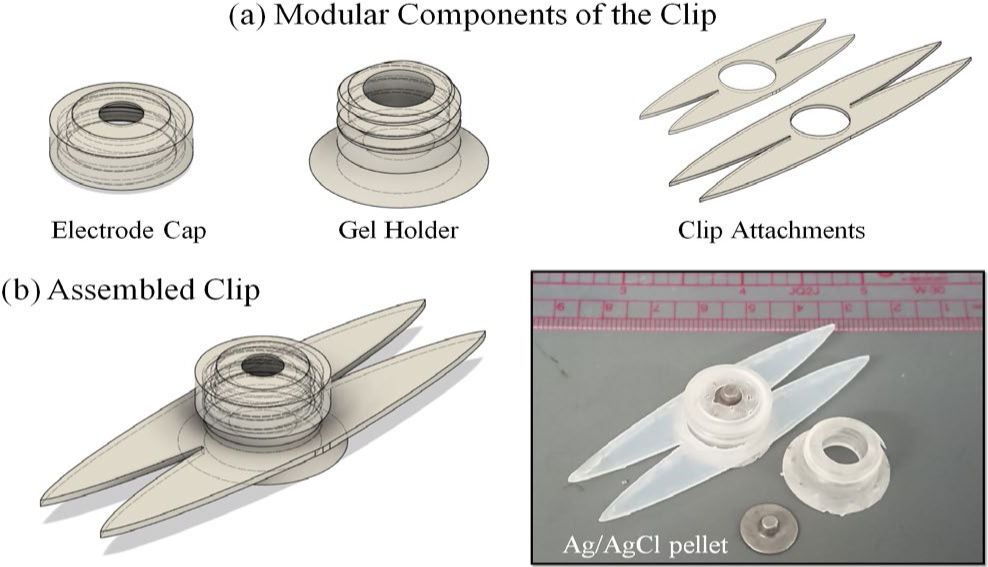
Components of the *Sevo* electrode prototype. (a) CAD drawings of the modular components of the electrode and (b) CAD drawing of the assembled clip shown alongside a photograph of the 3D printed prototype

## III. Methods

### A. Participants

Eight African-American participants from the ages of 18-24 with 4A-4C (Appendix A) hair types participated in this experiment.

### B. Electrode Design and Prototyping

Our design for the electrode-bearing hair clip (Fig. 3) is similar to that of a hair barrette in that the quantity and thickness of the patients’ hair stabilizes and holds the electrode firm against the scalp, thereby reducing impedance and improving the signal quality and signal-to-noise ratio. *In this sense, our electrode-bearing hair clip is able to turn what is conventionally viewed as a hurdle in EEG (i.e., African hair) into an advantage: it harnesses cornrowing, traditional in many cultures of African descendants, to improve scalp contact*. The design can be made in multiple sizes, such that the electrodes can be placed in different hair configurations as long as the housed electrode can make direct contact with the scalp through the hair. The parts of the clip are modular, allowing for easy resizing and cleaning between uses. In our early work, we found that braiding hair in a way that exposes the desired regions of the scalp makes electrode to scalp contact better. Combining that finding with this electrode design takes advantage of the mechanical properties of coarse, curly hair.

The base of the electrode was 3D printed in a resin-based printer of resolution 0.1mm (Formlabs, Somerville, MA, USA), and cured in an oven at 50°C while being exposed to UV light. The sides were laser-cut from polyethylene.

### C. Setup

Participants had one side of their head cornrowed in accordance with our developed method of cornrowing [5] to expose locations consistent with the 10-20 arrangement of electrodes, and one side of their head was left unbraided for fair comparison (see Figure 2). Using the 10-20 system of electrode placement, participants had gold cup electrodes (industry standard for EEG) placed at locations [7] F3, F4, FP3, FP4, P3, P4, C3, C4, O1, and O2, going along the scalp down both the braided and unbraided sides of the head. Participants also had *Sevo* electrodes placed at locations F4, FP4, P4, C4, and O2, going along the scalp down the braided side of the head. This variation in electrode placement was introduced to include the frontal, parietal, and occipital lobes of the brain, as well as test multiple orientations of the electrodes. All electrodes were applied with standard skin abrasion and conductive 10-20 EEG paste to increase the signal strength. The reference electrode was a Covidien Kendall surface electrode placed on a cleaned and abraded location on the mastoid, behind the ear.

### D. Process

The impedance of each electrode was measured using the Intan RHD2000 amplifier used with the Intan Recording Controller. The acquisition system used a sampling rate of 4kS/s, a notch filter at 60Hz. The electrode-skin impedance has direct influence on quality/SNR of EEG recordings [11]. Electrode impedance was measured in 5-minute time intervals for a total of 40 minutes at 30 Hz and 1000 Hz.

## IV. Results

Our experiment focused primarily on two aspects: how a) the braiding and b) the type of electrode affect the measured impedance. Overall, the impedance of our electrodes, combined with our cornrowing technique, is >10x lower than that of the clinical standard gold cups on unbraided hair. We also observed that over time, the impedance stayed relatively constant for all electrodes. The magnitude of impedance reduces as we increased the characterization frequency, as expected because capacitive electrode-skin impedance reduces with frequency. The frequency values chosen were 30 Hz (beta/gamma bands in EEG signals), and 1 kHz (the industry standard for characterizing the impedance of systems with mechanical/physical contacts). Evaluating impedance at two frequencies also enables recovering capacitive and resistive values separately if needed. Table I shows the highest values of impedance measured across time, averaged among all the electrodes we tested of a particular type/braiding pattern. Braiding the hair, along with our Sevo electrodes, performed about 20-30x better (Fig. 5) than the current practice of unbraided hair with gold cup electrodes.

**TABLE I.**
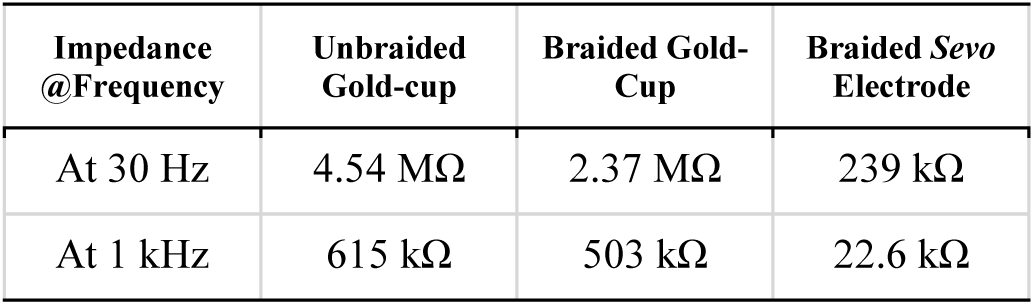
Highest Values of Impedance

**Fig. 4:**
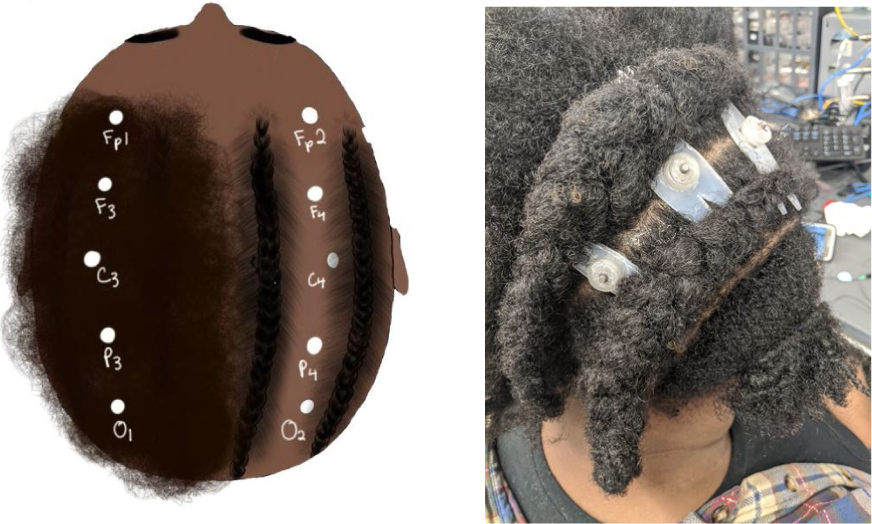
Our experimental protocol setup: The right half of each participant’s hair was braided, while the left half remained unbraided. Every labeled point bore a gold cup electrode, while the braided points bore a *Sevo* electrode as well. *(left)* Schematic diagram of electrode placements; *(right)* Photograph of *Sevo* electrodes placed on a braided participant.

**Fig. 5.**
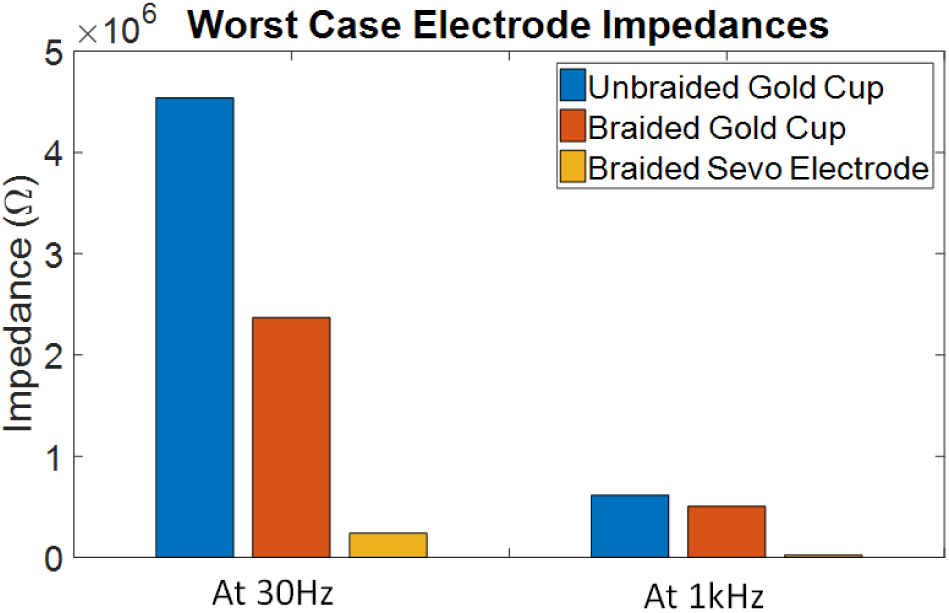
A plot of the measured impedances, recorded at their highest average value across time. The y axis is linear in scale. Braiding and using *Sevo* electrodes result in lower electrode-skin impedance.

Signal amplitude can be maintained by ensuring that electrode-skin impedance is less than the input impedance of the amplifier it connects to, which, for modern biopotential sensing amplifiers (e.g. TI chips) is ~500 MΩ [12]. Using our braiding technique, combined with the *Sevo* electrodes, the impedance (at low and high frequencies) is indeed negligible compared to 500 MΩ. Nevertheless, classical EEG systems advocate for 10-50kΩ resistance, and thus further advances are needed if lower-quality amplifiers are in use.

## V. Conclusions and Discussion

Our method of braiding the hair, combined with novel *Sevo* electrodes, preserves the integrity of the signal by using the properties of coarse and curly hair. Understanding the cultural significance and the unique structure of hair in individuals of African descent allows for a better patient experience as well as improved signal quality. This problem impacts a significant fraction of patients, as there are people of African descent in all parts of the world (including Afro-Latino and Afro-Caribbean, continental African, and African American populations).

Our system uses the properties of coarse and curly hair as a strength rather than a hindrance. For example, the process of cornrowing works better on individuals of Type 3 and Type 4 hair because of its relative coarseness. This coarseness prevents hair of these types from slipping out of the cornrows. Cornrows can be left in for up to eight weeks, making them easily usable for long-term measurement. Our solution is a beneficial technique for patients with seizures, where multiple measurements need to be made over extended periods of time. Because cornrows prevent hair from becoming the main source of impedance, our system leverages the properties of the hair that are generally considered a handicap in EEG data collection. Future work will examine this by observing our system with individuals of other hair types.

Typically, individuals of African descent are told to wash and straighten their hair without applying hair products, which can affect data quality. Depending on a person’s hair texture, this procedure may not produce desired results: due to the spring like nature of Type 4 hair (Appendix A), it is difficult to manage it without the use of hair products designed to tame curls. If any moisture is introduced by way of humidity, gel, or sweat, the hair begins a process colloquially known as “shrinking,” in which the hair reverts to its original curl pattern [13]. This can cause the technician to have to frequently re-apply the electrodes, or apply more pressure on them. The thicker and curlier a person’s hair is, the more prone their hair is to reverting back to curls and affecting the quality of data collection.

The current process of conducting EEG tests on patients with coarse, curly hair makes testing inconvenient for those individuals. This means that in neuroscientific and clinical research, fewer participants of African descent volunteer, and, as noted in the introduction (using [6]), post-recording, data from African descendants might be discarded unintentionally (simply because it had a lower quality to begin with) which could potentially be skewing research data, outcomes, neuroscientific understanding, and eventually, policy decisions. This also means that African descendants that do not have access to this core technology receive poorer quality of diagnosis for many disorders, or have an uncomfortable and/or emotionally taxing experience. This effect of the lack of understanding for hair type in the traditional method of conducting EEG tests deserves further and urgent attention from the neuroscience and engineering communities. That way, EEG can be more inclusive, without biasing against large sectors of the population.

## Acknowledgements

We thank Praveen Venkatesh for helpful discussions, and Yaramo Dione, Evangeline Mensah-Agyekum, and Sarah Korssa for cornrowing our participants.

## Appendix A: Hair Types

Hair for individuals of African descent is very diverse in texture. There are four subcategories to describe the curl pattern of curly hair for those of African descent. The subcategories of interest to our proposed problem are “Type 3” hair, which is categorized as looser, bouncier curls; and “Type 4” hair, which has tightly coiled, coarser, kinkier curls. Type 3 hair is easier to straighten without products than Type 4 hair, which is of our central interest and is more common amongst African descendants. Type 1 and Type 2 are straighter than Type 3 and Type 4 hair [8]. The radius of the curls is further classified by lettering each type, with “A” being larger curls, and “C” being very tightly coiled small curls. While our focus is on Type 4 hair due to the uniqueness of its texture, our method also works with Type 3 hair.

Fig. 6 of [8] illustrates the classification of hair curliness on a fine-grained scale, which further classifies hair type. The hair types of six different groups of ethnic origin. The results of this classification showed that people of African, African American, and Caribbean descent had much kinkier, curlier hair than those of Caucasian, Asian, and Brazilian descent.

## Footnotes

† Quoting Prof. Sarah Haigh (Psychology, Univ. of Nevada-Reno), “*Coarse and curly hair has a tendency to push against the cap, reducing the amount of contact between the electrodes and the scalp. We try to compensateby adding more gel to connect the electrode to the head, but this increases the likelihood of bridging electrodes. I run a lot of EEG studies on visual processing and hair is densest over the visual cortex. What often ends up happening is that we collect as much data as possible, then during preprocessing, the occasional dataset gets rejected from analysis for having too many noisy signals. Often, it turns out that those datasets are from those with coarse and curly hair - we have accidentally biased our own study by throwing out data with poor connections to the head!*” [6].

2 *Sevo* is the Haitian Creole word for “brain.” The name is motivated by the large fraction of African descendants in Haiti.

